# Simulating extracted connectomes

**DOI:** 10.1101/177113

**Authors:** Jonathan Gornet, Louis K. Scheffer

## Abstract

Connectomes derived from volume EM imaging of the brain can generate detailed physical models of every neuron, and simulators such as NEURON or GENESIS are designed to work with such models. In principal, combining these technologies, plus transmitter and channel models, should allow detailed and accurate simulation of real neural circuits. Here we experiment with this combination, using a well-studied system (motion detection in *Drosophila*). Since simulation requires both the physical geometry (which we have) and the models of the synapses (which are not currently available), we built approximate synapses corresponding to their known and estimated function. Once we did so, we reproduced direction selectivity in T4 cells, one of the main functions of this neural circuit. This verified the basic functionality of both extraction and simulations, and provided a biologically relevant computation we could use in further experiments. We then compared models with different degrees of physical realism, from full detailed models down to models consisting of a single node, to examine the tradeoff of simulation resources required versus accuracy achieved.

Our results show that much simpler models may be adequate, at least in the case of medulla neurons in *Drosophila*. Such models can be easily derived from fully detailed models, and result in simulations that are much smaller, much faster, and accurate enough for many purposes. Biologically, we show that a lumped neuron model reproduces the main motion detector operation, confirming the result of Gruntman[1], that dendritic compution is not required for this function.

## INTRODUCTION

Connectomes, maps of biological neural networks in a computer, are derived from volume imaging of the brain and include very detailed physical models of each neuron, such as the portion of an extracted neuron shown in Fig. 1(a). Simulators such as NEURON[2] or GENESIS[3] are explicitly designed to work with physical models, and compute results that depend on physical parameters. Driving a simulator such as NEURON with the output of EM reconstruction should therefore be able to reproduce the operation of biological circuits. However, straightforward attempts to do this run into several obstacles.

**FIG. 1.**
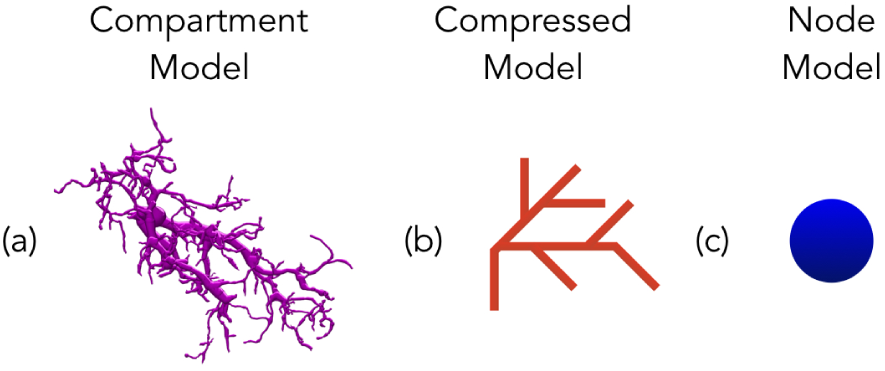
Panel (a) shows a fully detailed model of a portion of a T4 neuron, in the medulla of *Drosophila* (http://emanalysis.janelia.org/SharkViewer.php). Panel (b) shows (symbolically) a much simplified node model that preserves the branches but deletes intermediate nodes. Panel (c) shows the model reduced to a single node.

First, in addition to geometry, concrete numerical models of synapse operation are required. EM reconstruction can give synapse locations, but does not tell how they operate (or in *Drosophila*, even the sign-in-hibitory and excitory synapses look the same). Second, the use of detailed physical models brings additional concerns. The sheer number of nodes, plus the wide range of time constants between short and long segments, creates systems of equations that are hard to solve efficiently with numerical techniques. This not a problem unique to biological systems-circuit networks extracted from integrated circuits share the same concerns, and explicit techniques to avoid this problem have been used[4].

We investigated these problems by using the results from EM reconstruction to drive the simulator NEURON, to try to reproduce a known circuit operation-motion selectivity of the T4 cells in the medulla of *Drosophila*. We ran directly into the problems described above. First, we could not find in the literature detailed models for the graded synapses found in this circuit. Therefore we created analytic synapse models, tuned to get approximately the responses shown in the literature. Next, we found that if we used the fully detailed geometrical models, then the run times of the simulator were prohibitive. To proceed, we had to reduce the geometrical complexity of the extracted neurons. On a positive note, once we added plausible synapse models to our simplified geometrical models, we were able to reproduce major portions of the known network function.

To examine the tradeoff of geometrical complexity versus accuracy, we compared fully detailed simulations with several simpler models. These included both a simpler branched model and a model with a single lumped node. These simplified models are much smaller, much faster to simulate, and give nearly the same results for the neurons we consider here.

## EXPERIMENTAL DESIGN

For this experiment, we chose a portion of the visual pathway of the *Drosophila* fly brain, since it has both a detailed connectome[5], and a wide variety of detailed experimental and theoretical data. In particular we decided to try to reproduce the motion selectivity of the T4 cells. These cells react strongly to motion in the sensitive direction, and less strongly to other stimuli, including motion in the opposing direction, motion at right angles to the sensitive direction, or a uniform flash across the visual field. The T4 circuit is complex, with at least 8 differing cell types providing input, and the operation is still not fully understood[6][7][8][9].

The physical structure of each neuron in the network was imported as an SWC file, generated by the reconstruction of Takemura, et al.[5]. This network contains the full connectivity of the columnar cells of a 7 column core, and partial reconstructions of the cells one column further away.

Of the many pathways to T4, we chose the major ones starting with L1, based on synapse count. These include the excitory pathways L1 *→* Mi1, L1 *→* Tm3, Mi1 *→* Tm3, L1 *→* Mi4, L1 *→* Mi9, Mi1 *→* T4, Tm3 *→* T4, and the inhibitory pathways Mi4 *→* T4 and Mi9 *→* T4. We used all cells of these types that were contained in the 7 column reconstruction, resulting in 187 cells in our simulations.

For a stimulus, we applied an externally generated current to the L1 cells to simulate their excitation in the lamina, which was not included in the reconstruction we used. (In theory this could be added, since reconstructions exist for the lamina as well[10].) We simulate time varying visual patterns delays by adjusting the onset of each L1 stimulation (Fig 2(a))

The lamina L1 cells were stimulated in 5 patterns-a moving edge in each of the cardinal directions, and a full field flash. Since the L1 pathway, and the T4 cells, are thought to mediate the ‘ON’ response, we simulated a dark to light transition at each column. To create a moving edge, current was injected into different columns of the lamina with different delays-see Fig 2(a). We used an added delay of 100 ms per column. Since each column subtends about 5*°*, this corresponds to an edge moving at 50*°* per second, near the peak response speed for T4 cells.

For the L1 cells, whose inputs are not included, we injected current in a pattern that gave membrane potentials that approximate the response to a step function specified in the literature[11]. We settled on a step function of current, with 0.02 nA for 10 ms, 0 nA for 5 ms, and 0.0002 nA for 200 ms, as shown in Fig. 2(b). For the downstream synapses, since detailed models are not available, we constructed simplified models where the conductance of the membrane of the target is a function of the source synapse’s voltage level. We then adjusted these functions to get roughly the observed responses in the simple case of a step input to the cells concerned, where this was known. All cell models are graded response, not spiking.

**FIG. 2.**
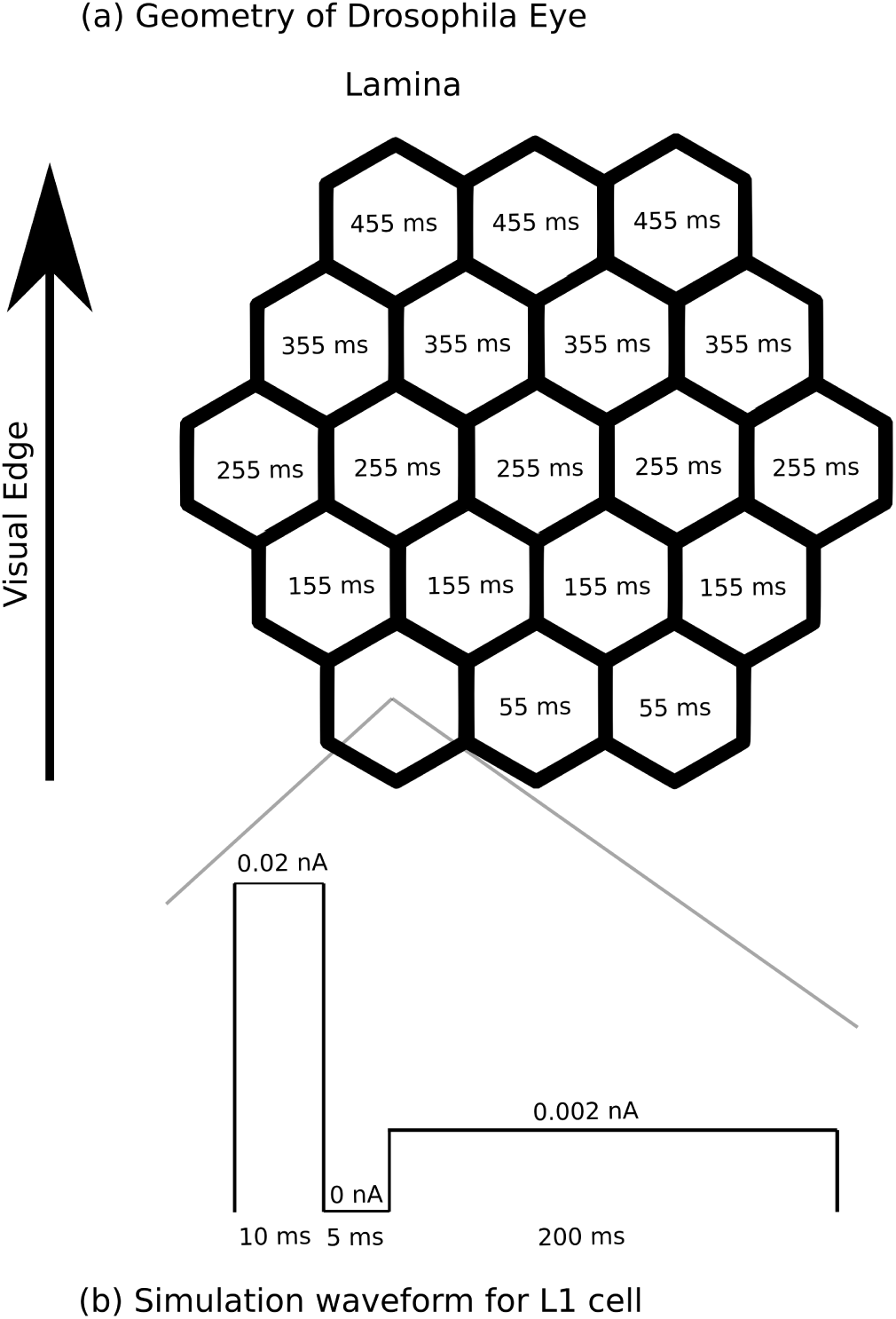
The construction of the lamina inputs in the *in silico* experiment. Panel (a) shows the hexagonal lattice of the fly’s eye. The numbers show, for the case of upward motion, when the stimulus for that column is applied. Similar patterns apply to the other directions of motion. In the case of a full-field flash, the stimulus is applied to all L1 cells simultaneously. Panel (b): the input that was applied to the L1 cells to get an approximation of the known response to a leading edge.

We then measured the response of each of the four subtypes of T4 cells to each of the 5 input patterns. What we hoped to show is that we could reproduce the main response property of these motion sensitive cells. Each should respond most strongly to an edge moving in its preferred direction, and less strongly (if at all) to motion in other directions or full-field simultaneous change in illumination. We did not try to match the measured T4 response over a range of conditions, as that is a research project in its own right[1], but instead just show different responses for different directions. This is sufficient for many uses of local motion detection, since opposing motion detectors are thought to be differenced in the lobular plate[12].

The expected mechanism for our responses is a Barlow-Levick motion detector[13], where excitation happens before inhibition in the preferred direction, inhibition occurs first in the non-preferred direction, and inhibition and excitation occur simultaneously in the cross-direction and uniform flash cases. The full operation of T4 is known to be more complex[14], but the subset of neurons we chose should support this computation.

In addition to our estimated synapse function, there are several other limitations to this experiment. The simulation includes only some of the columnar cells, and no non-columnar cells. Gap junctions were not included as they were not readily identifiable in the EM images used for neuron reconstruction, and hence not included in the connectome. Also only direct chemical synapses are used potentially longer range interactions such as neuromodulators are not modelled.

## MODELS OF NEURONS

The full compartment model neurons were generated from EM reconstructions. They are specified as a series of segments, with 3D endpoints for each and a radius at each end. This allows for the specification of neurons with a full branching physical structure. The extracted structure makes no distinction between axons and dendrites. This distinction is not relevant for insect neurons, whose neurites typically combine both inputs and outputs on the same branch. It also does not matter computationally-in NEURON, each neurite is conventionally named as an axon or dendrite, but computationally this is not significant.

In the absence of detailed biochemistry, each neurite was modeled as a passive leaky structure. This model has a constant conductance to a resting potential, plus variable leaky conductances (representing ion channels at synapses) that pull the voltage up or down. This model was chosen since this part of the insect brain consists of nonspiking neurons. The leak equation is a specified as:

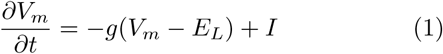

where *g*, a conductance, tells how strongly the membrane potential *V*_*m*_ is pulled towards the resting potential *E*_*L*_ (tyically -65 mV). In addition there may be a current *I* that flows into the neuron. In our simulations this is only used to excite the L1 neurons.

In NEURON, each neuron consists of cylindrical branches (the NEURON literature typically calls these soma, axons, and dendrites, though this distinction is not maintained in the fly). Each branch is a cylinder with geometric values of radius and length, obtained in our case from the SWC files. Finally, each branch is further divided into segments (compartments), where a soma typically has one segment and the dendrites have *n* segments, where the *n* is determined by the time constants of the branch geometry and the accuracy desired. (See section 5.7, “Choosing a spatial grid”, in *The Neuron Book*.)

The geometric values as specified in NEURON add membrane capacitance *C* and cytoplasmic resistance *R* to each neuron. This generates a model as shown in Fig. 3.

**FIG. 3.**
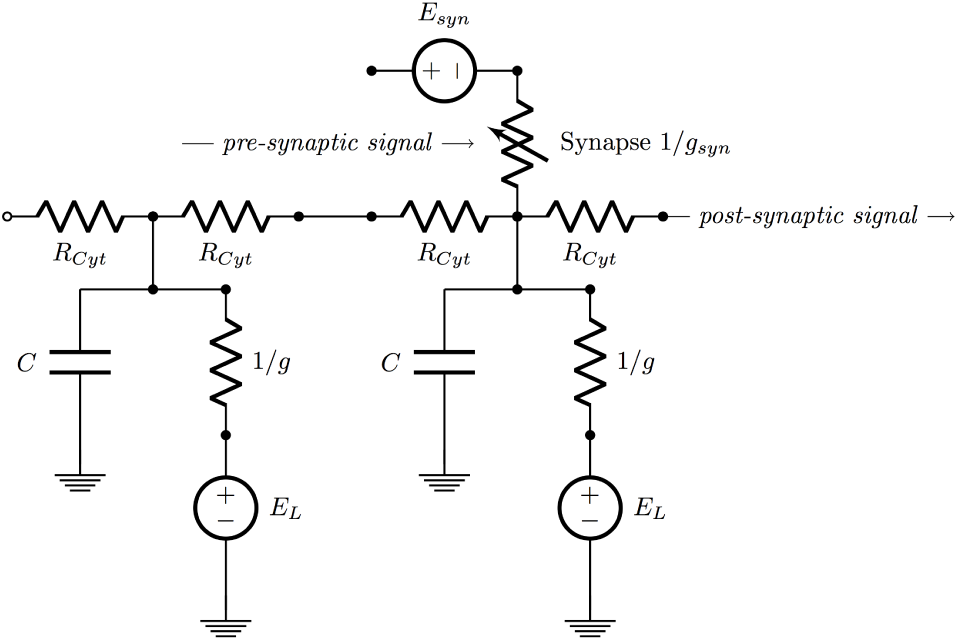
Circuit model for a compartmentalized, passive, leaky neuron.

### Adding Synapses

The synaptic connection between different neurons is a type of convolving synapse, with a synaptic depression term based on postsynaptic neuron membrane potential. This was not designed as an approximation of a biophysical molecular model, but rather intended to roughly reproduce the observed waveforms when combined with our passive neurite model. The equations we used are:

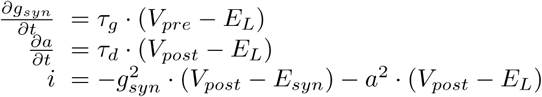

When a signal propagates through the synapse from the presynaptic neuron, the postsynaptic neuron membrane potential (mV) is pulled towards *E*_*syn*_, which is 0 for an excitory synapse and -80 mv for an inhibitory synapse. When the postsynaptic neuron membrane potential is high (low) enough, the synaptic depression term *a* grows slowly and starts to inhibit the signal propagating through the synapse[15].

When connecting neurons, the neural reconstruction specifies only the XYZ coordinates of each preand postsynaptic partner, separately from the SWC files that define the neuron geometry. Therefore connections were based on finding the two closest branches (one from the presynaptic neuron, and the other from the postsynaptic neuron) and then creating a synaptic connection between the two branches.

The types of these synapses (inhibitory vs. excitatory) were based on the findings from the *Drosophila* motion detection visual system [5]. The excitatory neurons used a *E*_*syn*_ leak value of 0 mv, while the inhibitory neurons had a *E*_*syn*_ leak value of -80 mv. Our synapses from pre-to post-synaptic neuron also required the parameters *τ*_*g*_ and *τ*_*d*_. We estimated these by approximating the membrane potentials of the experimental data [9] and the simulation results of the compartment models.

The equations in the network were solved using implicit Euler iterations, the default numerical method in NEURON. This method has first order accuracy in (Δ*t*), and computation time that in general is *O*(*N* ^3^), but is only *O(N)* in NEURON since it forbids loops[16].We recorded, for each neuron, the membrane potential as a function of time.

### Can a simpler model be used?

The models generated by EM analysis are very complex (for example, a typical Tm3 neuron has 2200 segments in the medulla), and it is not clear that the full level of physical detail is required. It is possible that simpler models may be faster to simulate, easier to understand, and still sufficiently accurate.

### Theoretical reasons simplified models might work

For sufficiently small neurons, theory suggests that membrane potential will be nearly equal throughout the cell, and simulating these potentials with simplified models will not sacrifice much accuracy.

We use a simple RC model to estimate time constants. The Elmore delay[17] *d* of a cylinder of diameter *D*, length *L*, resistivity *ρ*, and membrane capacitance *C*_*m*_, is

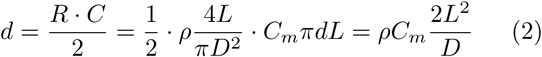

For the neurons we consider here, a thin branch might have a diameter *D* of 100 nm or 10^*-*7^m, and maximum length *L* of 50 *μ*m. Using typical values *ρ* = 1 ohm m, and *C*_*m*_ = 10^*-*2^ F/m^2^, we get a delay of 0.5 millisec. For comparison, we would expect the fastest response of this neuron to be about an order of magnitude slower, at about 5 millisec, based on a 1000*°* per second maximum turn rate of the fly and a 5*°* field of view of an ommitidium. We would therefore expect that the neuron will be largely isopotential and the voltage will not vary much from point to point in the neuron.

The brain of *Drosophila* is divided into many smaller compartments[18][19][20]. From the arguments above, it is likely that most neurons are almost isopotential within each compartment, but at least for fast signals, can have significantly different voltages in different compartments. Since delays depend on *L*^2^*/D*, not on either dimension alone, it is possible that this condition could hold in larger flies as well. The significance of this, if any, is unknown.

### Creating simplified models

In our first simulations, the aim was to make sure that the network was as close to the biological network as possible. Each cell in this network was typically composed of thousands of segments. Then, based on the analysis above, and experiments with a single neuron (where runtime was not prohibitive), we created simplified models with differing degrees of abstraction. For our first reduced model, we combined adjacent segments from the SWC file if there was no additional segment connecting to the junction between them. This model still has the branched structure of the original, but with fewer segments. Then for the simplest model, we compressed each neuron into a single node. In this model there is only one potential for the entire neuron, and so no possibility of differing computation in differing dendrites. The differences in the structures of the models and the populations can be found in Fig. 1 and Fig. 4.

**FIG. 4.**
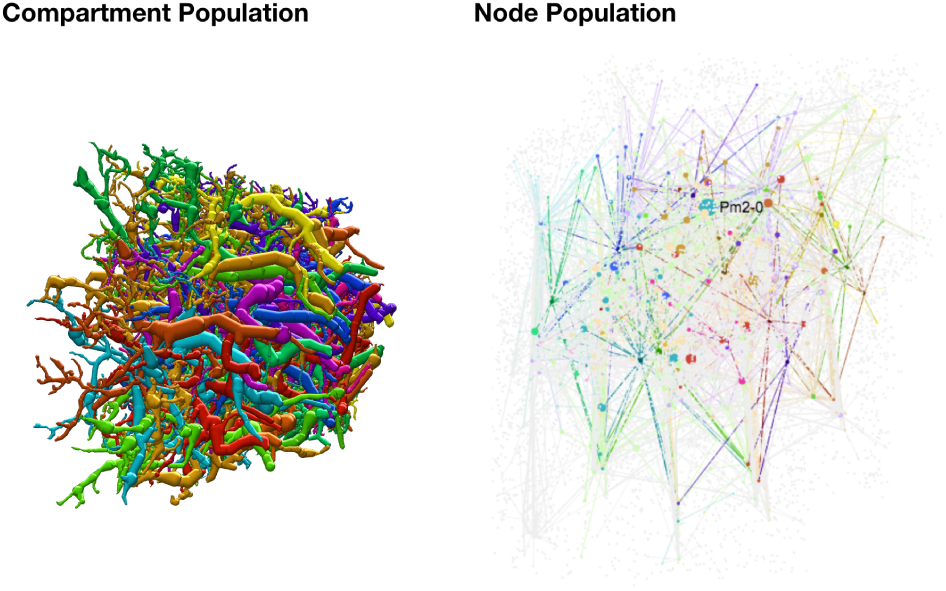
The differences of the population structure (http://emanalysis.janelia.org/gorgonian.php)

### Compressed node models

We reduced the model size by identifying every node that is not used by any synapse, and not a branch point. Each such node can be removed and its R and C values distributed to the adjacent nodes. In the case of DC response only, such a transformation is in principal exact, and no accuracy is lost. (Experimentally, of course, this is only true to double precision accuracy, but this inaccuracy is much smaller than other sources of error in our system). Computationally, this reduction can be done in time linear in the nodes deleted, by repeated use of the Y-Δ transform. After the resistance values (grounded and coupling) are computed, the capacitance values are distributed as proportional to the grounded (leak) conductances. This step is not exact but should be very close. This approach preserves the branching structure and hence allows for dendritic computation, but can reduce the size considerably. For one Tm3 neuron, for example, this simplification reduced the model size from 2214 nodes to 445 nodes, an 80% reduction.

To test the accuracy of this reduction, we ran detailed experiments with one neuron, with geometry and synapse locations from the reconstruction. The DC solutions were identical, as expected, within the limitations of computer arithmetic. The transient responses were evaluated using NEURON’s AlphaSynapse model, picking a subset of 20 synapses to fire simultaneously with a 5 ms time constant. Errors were largest where the synapses connect to the smallest diameter branches, so we picked the smallest branch to evaluate the error. Even so the error reached only a 0.2 mv difference between the full and the compressed model.

For the experiments where we included all 187 neurons, we used a similar but simpler procedure to generate the compressed models. We removed all interior, non-branch point nodes, and replaced each sequence of sections by a single section of average diameter. This was done for ease of experimentation, and we believe the error should still be small compared to the single-point approximation below. The more accurate (but harder to compute) model above could be used instead, if required.

### The single node model

The single-node model uses the same passive leaky neuron model that we used in the compartment model. Because the entire neuron is compressed into a single mode, the circuit diagram of the node model is quite different, as shown in Fig. 5.

**FIG. 5.**
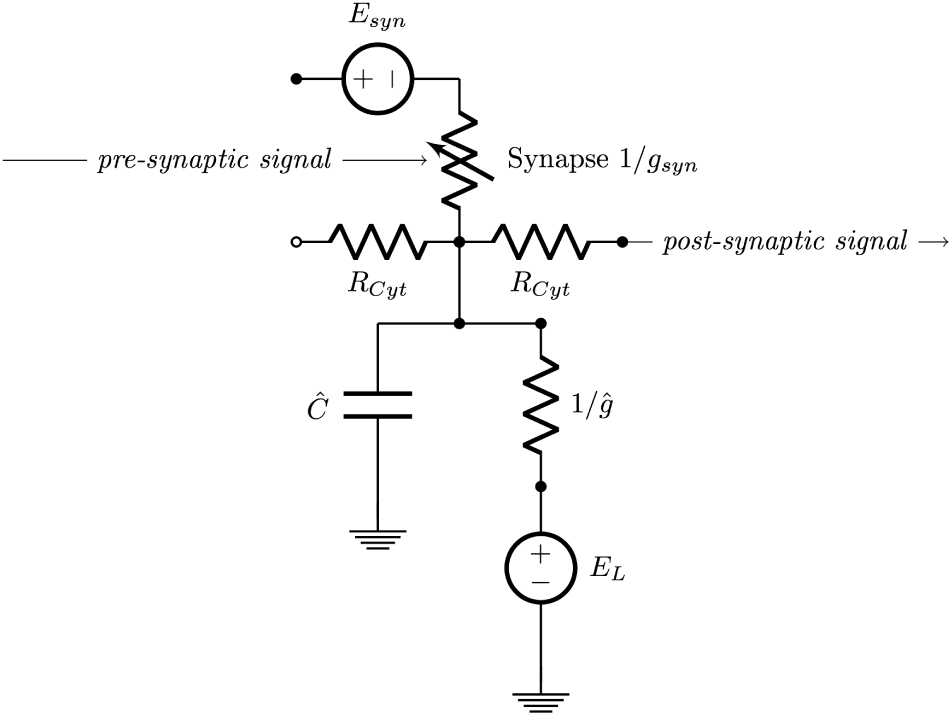
The equivalent circuit of the node neuron.

We wanted our single-node neurons to have the same total capacitance, and same leakage conductance, as the complex models simulated earlier. Since NEURON specifies these properties per unit area, to get the totals to agree we had to adjust the biophysical parameters of the single node neurons.

In the compartment model, both the capacitance *C* and the conductance *g* are summed over all the branches and segments of the neuron. Our single node model had a very different surface area, so we adjusted the new per-unit values to get the same totals. Specifically, we set a new capacitance *Ĉ*_*m*_ and and a new conductance ĝ. These parameters are set by first finding the total capacitance and conductance of the compartment model, by multiplying the per-area values of capacitance *C*_*m*_ and conductance *g* by the compartment’s total area *A*_*compartment*_ (which is calculated by a function within NEURON called area). Then these values were the divided by single node’s area to get the same total capacitance and conductance. The equation for each term follows:

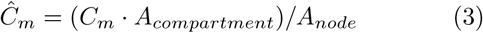

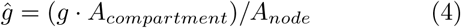

The node network used the same synapses with no changes, since the synapses have no geometry, and hence no area. The only difference is that the synapses no longer connect to the two closest dendrites (since the node model has no dendrites), but instead connect to the middle of each node.

## RUNNING THE SIMPLIFIED MODELS

We ran NEURON simulations to compare the different models, from the full compartment model to the most simplified point model. When comparing the full, reduced, and single-node model, we used one neuron simulated for 25 ms. When computing results on the whole network we used only the reduced and single-node models, as the full model was too slow. In this case each simulation ran for 1000 ms of simulated time for our network of 187 neurons. In each experiment, we recorded for each model the voltage as a function of time for all neurons. We then compared the results of the different simulations using external scripts written in MATLAB. In addition, we timed the simulations to measure the effect of the differing levels of detail.

Our results show that for a particular T4 neuron, the response is stronger in the preferred direction, compared to the opposite direction, the two orthogonal directions, or a uniform flash over the visual field. This result was obtained with either the reduced or single node model. See Fig. 6.

**FIG. 6.**
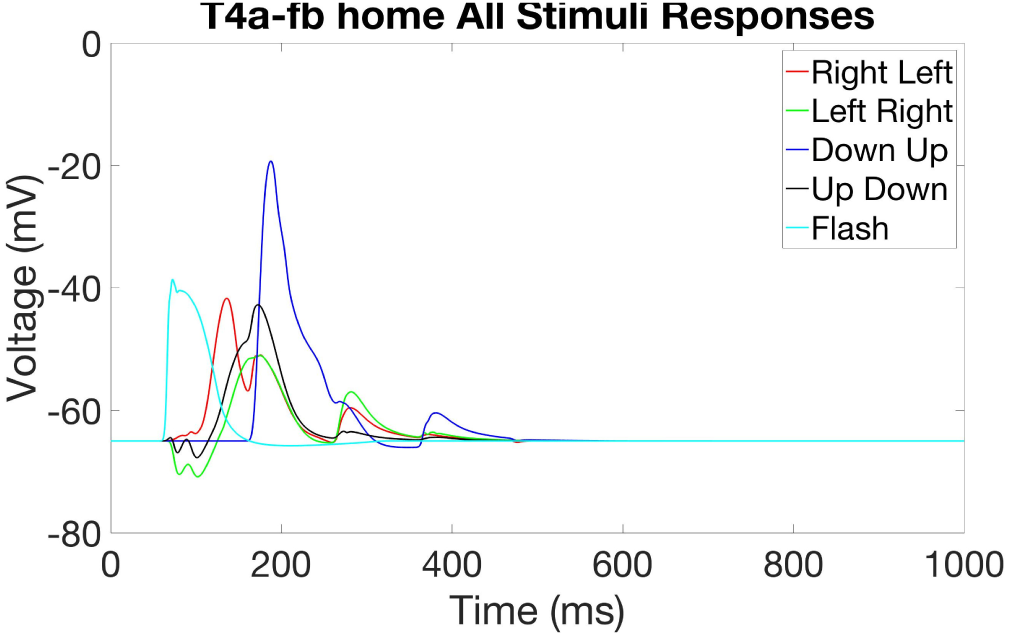
Response of one T4 cell (T4a-fb) in the home column to the five different test cases motion in the each of the four cardinal axes, and a full field brightening. As expected from the literature, there is a preferred direction for responses.

Similarly, the four different types of T4 cells, thought to encode a particular direction, do so as expected. Again this result was observed with both the reduced and single node models. See Fig 7.

**FIG. 7.**
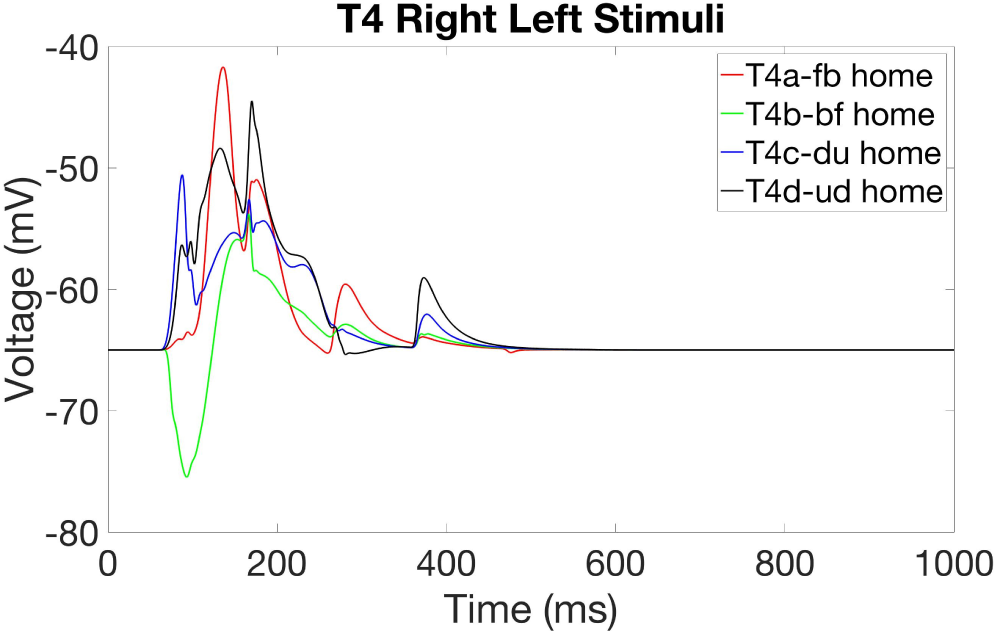
Response of the 4 different T4 sub-types to a right-to-left stimulus. The fb subtype gives the largest response, the opposite direction (bf) the smallest response, and the two cells with receptive fields at right angles (du and ud) give an intermediate response.

Comparing the different levels of detail, in Fig. 8, we show the recorded membrane potentials from each of the T4 cells (a neuron population in the motion detection circuit in the *Drosophila* visual pathway), for the reduced model and the simplest single-node model. The T4 cells were chosen for display since they are at the end of the visual pathway, and hence most likely to be affected by changes to the network structure. As seen in Fig. 8, the waveforms computed from the detailed model and the node model are very similar-by eye, they are almost indistinguishable. Fig. 9 shows the computed differences between the membrane potential of the compartment and the node model. Even in this case, with no consideration whatsoever of neuron shape, the voltages differ for only a few nodes and only by a few tenths of millvolts.

**FIG. 8.**
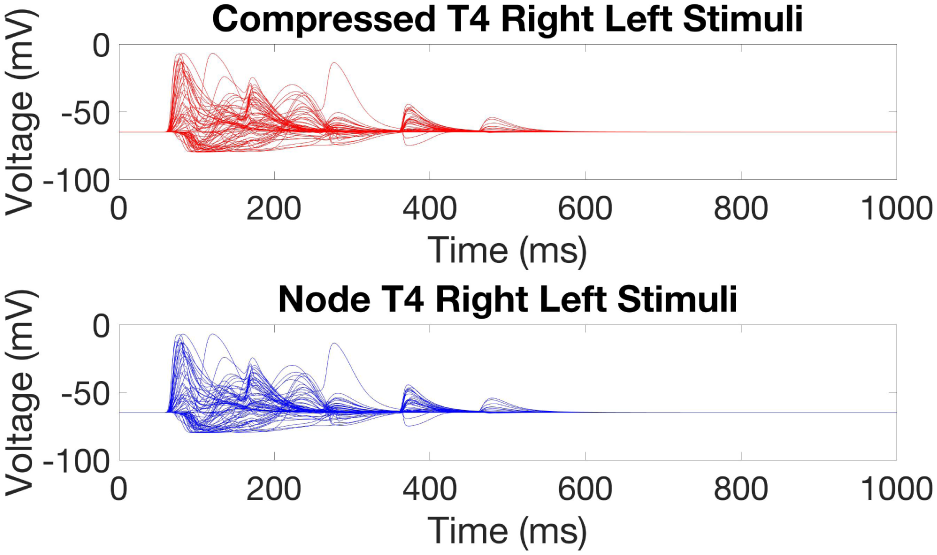
The membrane potential of the T4 neurons for both the reduced and single-node models.

**FIG. 9.**
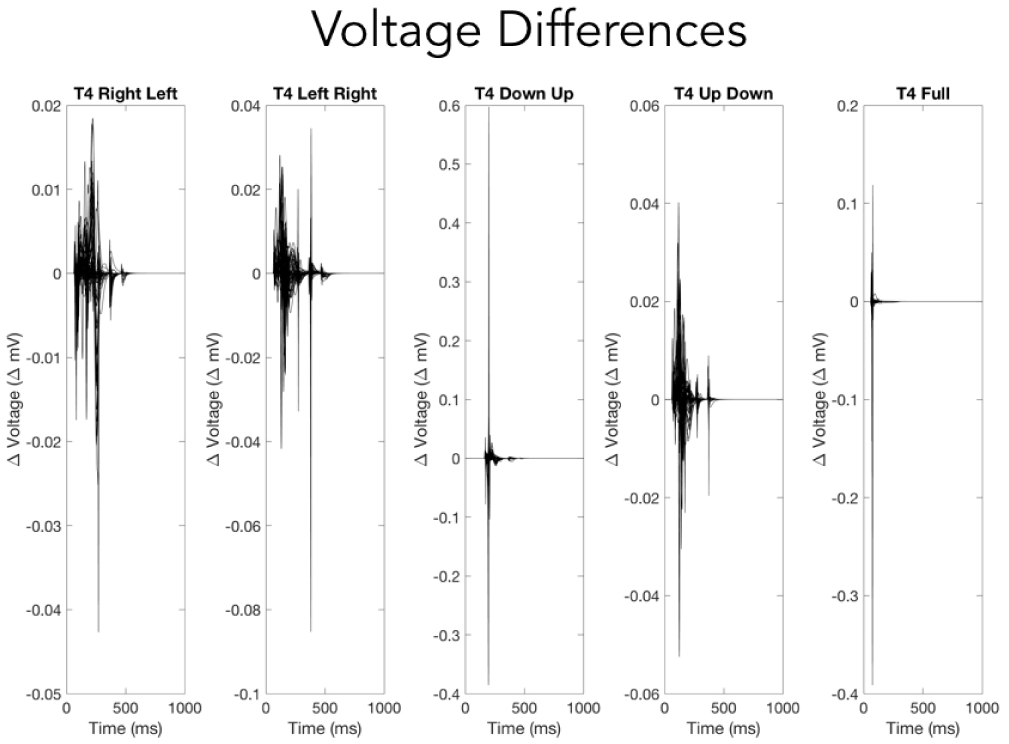
The differences between the T4 neuron membrane potential of the compartment and the node models.

### Simulation Times

To compare the run times of the simulations, we attempted to run 10 simulations of the full compartment model, the reduced model, and the node network. For simplification we ran just one case, where all the L1 neurons were stimulated at the beginning of the simulation. We ran 10 simulations for a simulated time period of 100 ms, 200 ms, etc for all models. We stopped the full compartment simulation before it completed, after one day of running time. Fig. 10 shows the differences in simulation times. The single-node model is at least 4 orders of magnitude faster than the full compartment model.

**FIG. 10.**
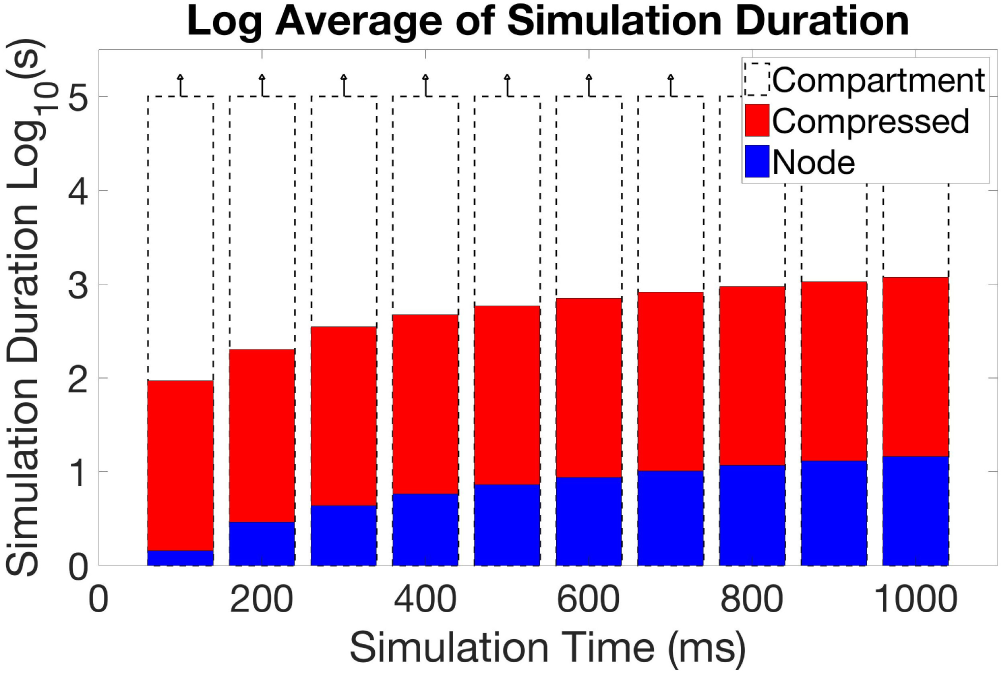
The run-time differences between simulation with the full model (200,000 nodes), the reduced model(28,000 nodes), and the node model (187 nodes). The full model times are underestimates since we killed the simulation after one day (roughly 10^5^ seconds).

### Comparing the node and detailed models

Based on the comparison of the compartment and the node model, there are two important takeaways. The first is that the efficiency of the simulations can be increased by using simplified models without sacrificing network function and dynamics. The second is that, at least in the medulla of *Drosophila*, the connectivity of a neuron is much more important than its detailed shape. Dendritic specific compution is not needed to explain the observed function.

Simulation experiments can require execution time ranging from from seconds to days, depending on the level of detail and the time span simulated. Using very detailed models, as obtained from EM reconstructions, while very accurate, contributes to these long run times. Here we show, at least for the medulla of *Drosophila*, it may be unnecessary to run simulation experiments on the fully detailed compartment models. Instead, it may be preferable to simplify the model, possibly as far as a single node model, as the simulations will run faster and the network dynamics are not sacrificed when moving between the two models. In neuroscience, in many cases form leads directly to function. But in our simulation experiments here, we find that the detailed form does not lead to function; it is rather other properties of the neuron that are important for function. In particular, the connectivity seems much more important than the shape.

## CONCLUSIONS AND FUTURE RESEARCH

It is possible to run detailed neural simulations using the fully detailed shape models obtained from EM reconstruction, but it is very slow. Models that are at least somewhat simplified are needed to get practical runtimes for circuits with a few hundred neurons.

These simulations, using plausible synapse models, can reproduce many experimentally determined aspects of neural computations. In particular, the main behaviors of the motion detectors in *Drosophila* can be reproduced from extracted neuron models and likely synapse signs.

A major limiting factor for neural simulation appears to be the lack of detailed synapse models. This is not completely clear, however, as other aspects of neural operation, such as gap junctions and neuromodulators, are also known to be missing from existing models.

It seems likely that within a single compartment of the *Drosophila* brain fully geometrically detailed models are not required. In many cases much simpler models, even single point models, can be used with only minimal loss of accuracy. In particular, our experiment with the T4 (the on-pathway motion detection circuitry of *Drosophila*), demonstrates that at least for this circuit, we can use point models instead of fully distributed models with only a minor loss of accuracy. Therefore dendritic compution is not needed to explain this computation, confirming the result of Gruntman *et al*.

For the future, reductions are shown in the paper will likely become more important, as much larger EM reconstructions are under way. This will presumably lead to desires to simulate much larger portions of the fly’s brain, including both more and larger cells. Several reasons suggest that reducing the neurons to a single point will not suffice for these larger circuits. They are both physically larger, with larger internal time constants, plus they will likely include spiking neurons with faster internal dynamics. For these larger and faster neurons we would expect that although they could not be reduced to a single point, reduced models will still be required to keep simulation times practical. Our research indicates this is feasible without undue accuracy penalty, but further experiments with networks of spiking neurons would surely be desirable.

